# The changing status of imperiled species in British Columbia over the last 15 years in the absence of a dedicated species at risk law

**DOI:** 10.1101/2025.09.30.679339

**Authors:** Peter R. Thompson, Meg Bjordal, Morgan L. Piczak, Sarah P. Otto

## Abstract

Gaps in biodiversity protection occur in Canada due to limited jurisdiction of federal species-at-risk laws and the absence of dedicated legislation in many provinces, including British Columbia (B.C.). While lacking legal protection, B.C. maintains Red, Blue, and Yellow Lists of threatened, special-concern, and secure species, respectively, using a NatureServe ranking system. We compiled historical data on species status in B.C. between 2008 and 2024 from the B.C. Conservation Data Centre. B.C. is home to 5,485 Yellow, 1,116 Blue, 491 Red-listed species. Changes in status over this time period were reported for 967 animal species, with an even split in uplistings (more imperiled) and downlistings (less imperiled). More status changes were reported for plants (2,902), mainly due to updated methodology leading to a lower risk status. Analysing the accompanying explanation for each status change revealed that most changes were non-genuine (e.g., new information, taxonomy, or methodology) rather than genuine (e.g., true changes in population size, range, or threats). Genuine improvements in the status of species in B.C. have been exceedingly rare. This analysis indicates that current laws and regulations have been insufficient to recover species at risk within B.C.

**Plain-language summary:** British Columbia is the most biodiverse province in Canada, yet 1,607 species are imperiled in the province. We analysed trends, finding a 16% rise in the number of species at risk since 2008. By analysing the provided explanations, we found that status improvements were mostly “non-genuine”, caused by changing methodology and/or new data. Strengthening legal protections is essential to prevent biodiversity loss in the province.

## INTRODUCTION

Legislation such as Canada’s *Species at Risk Act* (SARA) and the United States’ *Endangered Species Act* (ESA) provide crucial legal frameworks to prevent extinctions, support species recovery, and hold governments accountable to mitigative actions. These acts can be effective; for example, the *ESA* has contributed to the recovery of 291 species since its enactment in 1973, significantly reducing extinction rates across many species (Greenwald et al., 2019). In Canada, *SARA* applies primarily to federal (Crown) land, prompting many provinces and territories to implement their own complementary legislation (e.g., Ontario, Manitoba, Nova Scotia, New Brunswick). Several provinces lack dedicated species at risk laws, however, creating critical gaps in protection.

British Columbia (B.C.) is the most biodiverse province in Canada (CCA, 2020), yet it lacks a dedicated species-at-risk law (Gordon et al., 2024). While B.C.’s *Wildlife Act* (1996) has provided some protection against harm for species at risk, *only* four species have been listed since its inception: sea otter (*Enhydra lutris*), Vancouver Island marmot (*Marmota vancouverensis*), burrowing owl (*Athene cunicularia*), and American white pelican (*Pelecanus erythrorhynchos*). Other provincial legislation, such as the *Forest and Range Practices Act* (*FRPA*; 2002), provides limited and fragmented protection (FPB, 2023; OAG B.C., 2021; Province of B.C., 2023). For example, there is no legal requirement for recovery planning under *FRPA*, and the Act only applies to provincial lands under forestry or range tenure, therefore constraining its effectiveness in providing protection. The lack of dedicated legislation is particularly alarming given that 337 extant species have been found to be at risk of extinction in B.C. by the federal Committee on the Status of Endangered Wildlife in Canada (*COSEWIC*), with 256 receiving federal protection under *SARA* (i.e., on Schedule 1, Government of Canada, 2025). Most protections under *SARA* apply automatically only to the 1% of B.C. lands that are federal (Bollinger et al., 2020), with the province holding primary responsibility for species protection.

While B.C. does not have a legal framework to protect species, the province does assess the status of species through the B.C. Conservation Data Centre (CDC), a provincial government organization whose mission is to help conserve biodiversity by collecting and sharing information about wildlife and which is a member organization of NatureServe. Information for each species in the province, whether at risk or not, is posted online in the B.C. Species & Ecosystems Explorer (https://a100.gov.bc.ca/pub/eswp/). This information is gathered from a variety of sources, including museum collections, published and unpublished reports, expert consultants, and community science, and is subsequently screened by CDC staff. In addition, assessments conducted by COSEWIC are shared and can inform assessments within B.C. Each species or subspecies in B.C. with sufficient information is assigned a provincial status based on NatureServe ranks (“S Rank”, where “S” stands for “subnational”; Faber-Langendoen et al. 2012), ranging from S1 for critically endangered to S5 for secure. S Ranks also communicate the level of certainty associated with a species’ assessed level of risk; for example, a ranking of “S1S2” indicates that the available information on a species is not precise enough to determine whether it should be assigned to S1 or S2, and a ranking of “SU” (“U” for “unknown”) indicates even higher uncertainty. This S Rank status is then used to assign species to the colour-coded “B.C. List”. Species on the Red list are native species that are endangered, threatened, or extirpated and generally have an “S Rank” of S1 or S2, indicating they are imperiled. Species designated as Blue are native species of special concern and typically have an “S Rank” of S3, meaning they are considered vulnerable. Species listed as Yellow are not considered to be at risk and have more secure S Ranks (e.g., S4 or S5), reflecting more secure conservation status. The precise relationship between the S Rank and the B.C. List, accounting for uncertainty, differs slightly for plants and animals (e.g., an assignment of S2S3 is Red-listed for animals but Blue-listed for plants; see B.C. Conservation Data Centre, 2023). Currently, the B.C. CDC tracks over 24,000 species and subspecies in B.C., with over 10,000 assigned S Ranks.

While these ranks provide a useful snapshot of the conservation status of species in B.C. and offer a foundation for improving protections, the <u>B.C. Species & Ecosystems Explorer</u> database (B.C. Conservation Data Centre 2025) only displays present-day (“live”) species statuses and ranks, making it challenging to detect trends over time and the reasons underlying these trends. The B.C. CDC also publicly shares annual reports summarizing database updates, with brief explanations on status changes, as well as past snapshots of the database. The evidentiary basis for status assignments or changes is challenging to evaluate externally, as it is only briefly summarized in these reports and is based on an internal process (potentially consulting external experts) without peer review or detailed reporting. To address these challenges, we compiled archived datasets from 2008 to 2024 to create a static database and examine long-term trends in the statuses of species at risk over time. Here, we summarize changes in the number of Red-, Blue-, and Yellow-listed species over time, distinguishing between newly added species and status changes for already listed species. By analyzing explanations in the annual reports, we also evaluated what fraction of these status changes represented genuine improvements or declines in the status of species in B.C. We then compare changes to species status federally and provincially. Finally, we present four case studies that illustrate the changing status of species at risk in B.C.

## METHODS

### Trends over time in status

We accessed archived data containing historical information on species “S Ranks” and corresponding B.C. List assignments between 2008 and 2024 (provided by the B.C. CDC on February 5, 2025, now archived for 2011-2024 by the Government of British Columbia, 2025). Each year had a corresponding file (typically, a CSV file or Excel spreadsheet; in some years, animal and plant taxa were separated into two files) containing information on species (or subspecies / subpopulation) names and statuses. These datasets contained taxonomic information, provincial S Ranks, and federal assessment and listing information based on COSEWIC assessment reports and status under *SARA*. The datasets also indicated which species were native (i.e., up for consideration for Yellow, Blue, or Red listing) and non-native (and considered “Exotic”), as well as “Accidental” species that have been reported in B.C. only in rare, exceptional circumstances (e.g., Pinyon Jay *Gymnorhinus cyanocephalus*, which has only been recorded twice in the province; Toochin, 2024).

Merging these datasets together in a way that joined assessment histories for individual species was not trivial because the formats of historical datasets varied across time. A common cause of variation was taxonomic changes including splits (where a species or population was divided into two separate species or populations in a subsequent year), lumps (where multiple species and/or populations were combined into one in a subsequent year), or changes to English and/or scientific names. Fortunately, most datasets (excluding the years 2008 and 2009) assigned “element codes” to each species or population that were typically robust to name changes. We iteratively grouped species together by matching element codes from a previous year with those from the subsequent year, starting with 2023 and 2024 since those years had the most species in the database. For any element codes that did not match the subsequent year (e.g., element codes from the 2023 data that did not appear in 2024), we searched for any entries from the subsequent year with the same scientific name. If there was still not a match, we attempted to match species using old scientific and English names for each species (e.g., a value in the “old scientific name(s)” column in the 2024 dataset could be the scientific name that was in use in 2023). We tried the same process with English names, but since these were sometimes used for more than one species (particularly for invertebrates), we only matched with English names if the name only appeared once in the entire dataset. Any species that still did not have a matched entry for the subsequent year was evaluated manually and, if necessary, assigned a new element code.

Once we had processed and unified the datasets for each year, we assessed inter-annual trends in both S Ranks and B.C. List status. We also confirmed that the one-to-one matching relationship between S Ranks and listing colours used by the B.C. Conservation Data Centre (2025) applied for all species. In particular, we were interested in assessing the presence of species that were not Blue- or Red-listed despite S Ranks sufficient for Blue- or Red-listing, which we termed “ghost” species because the evidence used to assign an S rank leaves no trace in the B.C. List. We assessed trends in S Ranks, listing colours, and ghost species across taxonomic axes as provided by the input data.

We also compared species statuses at provincial and federal levels to determine the degree of concordance between these two assessments of extinction risk. We acknowledge that COSEWIC and the B.C. CDC may correctly assess a species to different levels of risk owing to the differing spatial extent between focus areas. The federal government also divides many species into “designatable units” representing discrete and evolutionarily significant populations, which may be treated with different listing statuses; in these cases, to be conservative, we compared B.C. List values with the designatable unit that was assigned the lowest level of risk by COSEWIC. For example, the Eulachon (*Thaleichthys pacificus*) is Blue-listed at the species level in B.C., but has three separate designatable units (assessed by COSEWIC as Endangered, Endangered, and Special Concern) federally, so we compared Blue (provincial) with Special Concern (federal) in this case.

### Changes in status

We downloaded information about changes to the name or status of animal species or non-animal species (i.e., vascular plants, lichens, macrofungi, and slime molds, herein termed “plants”, which form the majority of these taxa) from 2008 to 2024 from the B.C. CDC (https://www2.gov.bc.ca/gov/content/environment/plants-animals-ecosystems/conservation-data-centre/explore-cdc-data/conservation-data-centre-updates; accessed 6 March 2025). The data were amalgamated and standardized to obtain a consistent set of header names (see Supplemental 1).

To examine trends across all taxa, NatureServe S ranks were converted into numerical values based on the average value assessed, dropping any additional coding (e.g., a value of “2” was assigned to S2, S1S3, S2?, while a value of “1.5” was assigned to S1S2). Changes in certainty were also examined, with ranks including “?”, “SU”, or “Unknown” coded as less certain.

Among animal species, there were 162 changes in risk status on the B.C. List (Red, Blue, Yellow). In each of these cases, a text-based “rank change comment” was provided in the changes database, which we included in further analysis. In half of these cases (*n*=81), a “rank change reason” was also provided. We first scored the comments and reasons as to whether there was a true change to the status of the species (“genuine”), reflecting trends in population size or range, rather than “non-genuine” changes, which included reasons such as updated methodology, the inclusion of new information, altered search effort, or taxonomic changes (guided by IUCN standards, IUCN, 2025 ; assessed 23 March 2025). These definitions of “genuine” and “non-genuine” changes align with those used by NatureServe (2025, see also Teucher and Ramsay 2013). NatureServe (2025) notes that genuine changes can be difficult to determine and recommends to assessors that a genuine change be recorded only if “something has actually changed on the ground (or in the water) regarding the population/occurrence size, health, locations, trends for, or threats to, the element.” Examples of non-genuine rank changes include comments such as: “The rank change is due to a re-assessment of the original information, using current ranking methodology” or “New information. There has been an increase in range and number of occurrences due to increased survey efforts.” For each species with a change in listing status, three authors [MB,PRT,SO] independently interpreted whether the text in the “rank change comment” and “rank change reason” columns corresponded to a genuine or non-genuine change in rank, while blind to the species and status columns. Many comments regarding listing changes were ambiguous, often providing evidence for the current status but not explaining the reason for the status change. Consequently, interpretations varied across authors as to whether the change was genuine in 17.9% of *n*=162 cases (Table 1). We checked our evaluations against the animal databases for 2013, 2014, and 2019, when numerical codes were used for whether the change in status was genuine (*n*=25). When the data code indicated that the status change was non-genuine (*n*=22), we were unanimous that the change was non-genuine in 20 cases and varied in our interpretations in 2 cases (with 1 out of 3 of us scoring as genuine). When the data code corresponded to “genuine” (3 cases), we agreed in one case:

**Table 1:**
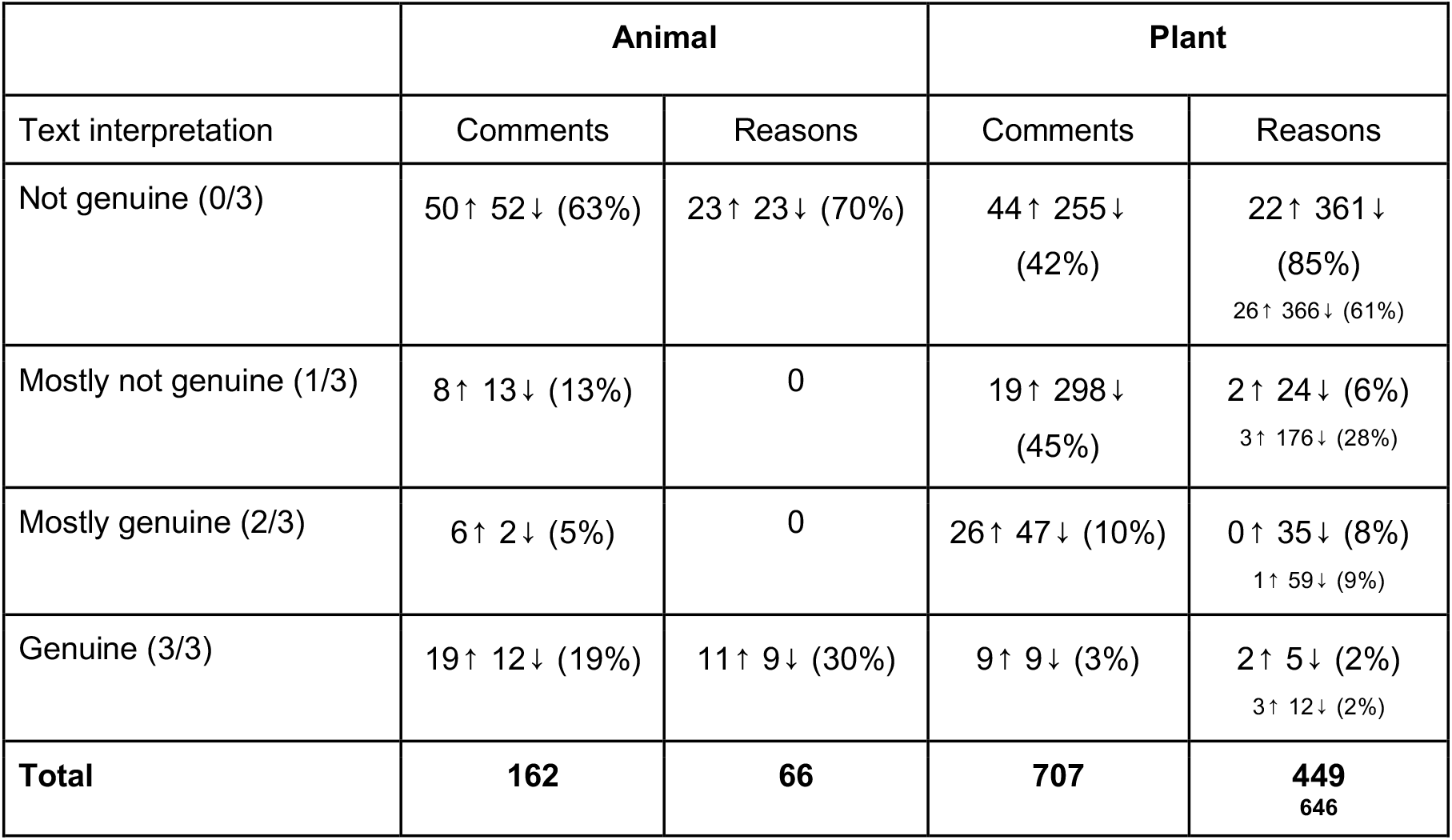
Genuine vs non-genuine changes in B.C. List status (Red, Blue, or Yellow). Comments (less reliable) and reasons (more reliable) were interpreted independently by three authors for species that became more endangered (↑ uplisted) or less endangered (↓ downlisted). Numbers in the first column refer to the fraction of evaluators that interpreted the text as a genuine reason for change. For plants, the reasons in large font are for 2019 onwards, while the small text below also includes the more ambiguous reasons given in 2018.

> *Hesperia colorado oregonia* (butterfly): “Western Branded Skipper habitat, including Garry oak and coastal sand ecosystems, are highly threatened and of the 16 historical sites known, only 4 remain extant.” [Ranked as genuine by 3/3 of us]

The other two species ranked as a genuine change in the database as deemed by the B.C. CDC:

1. *Recurvirostra americana* (bird): “This change in status is likely a combination of increased range and increased search effort. It is a distinct, showy bird and some of the places that American Avocet have been found are not particularly remote supporting that this is not just a case of increased effort.” [Not ranked as genuine by any evaluators]
2. *Dryobates villosus picoideus* (bird): “Numbers of Hairy Woodpeckers are confirmed to be stable to increasing. This also is the first time assessed with rank calculator as well as a full threat calculator confirming low threats.” [Ranked as genuine by 1/3 of evaluators]

We are thus cautious about our interpretations of the “comment” column for evaluating the nature of the change in status. In 2019, a “rank change reason” column was added that provided more standardized reporting (e.g., “Genuine change in status (time period uncertain)” or “New interpretation of same information”). For species with a status change between 2019-2024 on the colour-coded B.C. List (Table 1), our scores were unanimous for this “reason” column, perfectly matching the numerical score given in 2019 for genuine/non-genuine changes.

Among plant species, a text-based “rank change comment” was included for many of the early years. As with animal species, whether comments were genuine or not was often ambiguous, and interpretations varied amongst the three evaluators for 43.4% of cases (*n*=1,301). Starting in 2018, text-based “rank change reasons” were provided, which were interpreted more consistently among the three evaluators (20.6% varied in score, *n*=1,179). The reasons given in 2018 were particularly ambiguous (91.3% of our scores varied, *n*=196), but the reasons became standardized in 2019 and subsequent years (6.5% varied in score, *n*=983). Table 1 summarizes these evaluations for species whose status changed on the B.C. List (Red, Blue, Yellow).

We then evaluated the proportion of uplistings (more imperilled) and downlistings (less imperilled) that involved genuine versus non-genuine changes, using both the rank change comments (2008-2024) and reasons (restricted to 2019-2024), with the latter being more reliable.

### Statistical tests

To assess whether species whose status had improved or declined were more likely to reflect genuine versus non-genuine change, contingency tests were conducted in *Mathematica* using Fisher’s exact tests to account for categories with low numbers.

## RESULTS

### Trends in the Status of Species in B.C.

As of February 10, 2025, there were 24,191 species in B.C.’s Conservation Data Centre, of which 3,665 had not yet been assessed, 2,203 were not applicable to the listing process (either “Exotic” or “Accidental” species), and 7,758 had been assessed as “SU” (indicating high uncertainty in the species’ status in the province). This leaves 10,565 species that had been assessed by the provincial government, given an S rank, and so eligible to be placed on the B.C. List. Of these species, 5,485 have been listed as Yellow, 1,116 as Blue, 491 as Red, and the remaining 3,473 species lacked status on the B.C. List.

Of the 3,473 species that have a numeric NatureServe status rank but do not appear on the B.C. List, 1,012 warrant listing as species at risk in B.C. according to B.C. CDC criteria, with 868 on the Blue List and 144 on the Red List. Almost all of these “ghost” species (*n*=991; 97.9%) are arthropods, with the remainder being sponges (*n*=3), vertebrates (*n*=17), and one moss, with hundreds of these species having been unlisted since 2017. In 2022, the B.C. List status changed for many ghost species from “No Status” to “Not Reviewed”, suggesting that a lack of resources to verify assessments may be preventing these ghost species from having official status on the B.C. List. Other species that merit review include three bird species and one fungus that were assigned a “colour” (three Yellow, one Blue) when their S Rank indicates a more at-risk status (three Blue, one Red, respectively).

In 2008, the B.C. CDC database included dramatically fewer species (5,791 in total, 24.0% of the 24,191 species in the 2024 database). The 2008 database also included fewer ghost species (*n*=13, 0.2% of the dataset) and fewer unreviewed species (*n*=17, 0.3% of the dataset). Much of the growth in the database over the past 15 years reflects expansion to include a wider diversity of species. In 2008, the majority of species in the database were flowering plants (51.7%), but this proportion decreased to 13.5% in 2024 owing to the inclusion of many insects (making up 45.0% of the 2024 database, compared to 7.1% in 2008) and fungi (13.2% in 2024 versus 0.1% in 2008). The majority of newly added species were placed on the Yellow List or assigned No Status (Figure 1). For example, in 2017, 5,406 invertebrate species were added (mostly beetles), many of which were unknown in status (SU) or were “ghost” species with a numeric S ranking but “No Status” or “Not Reviewed” on the B.C. List.

**Figure 1:**
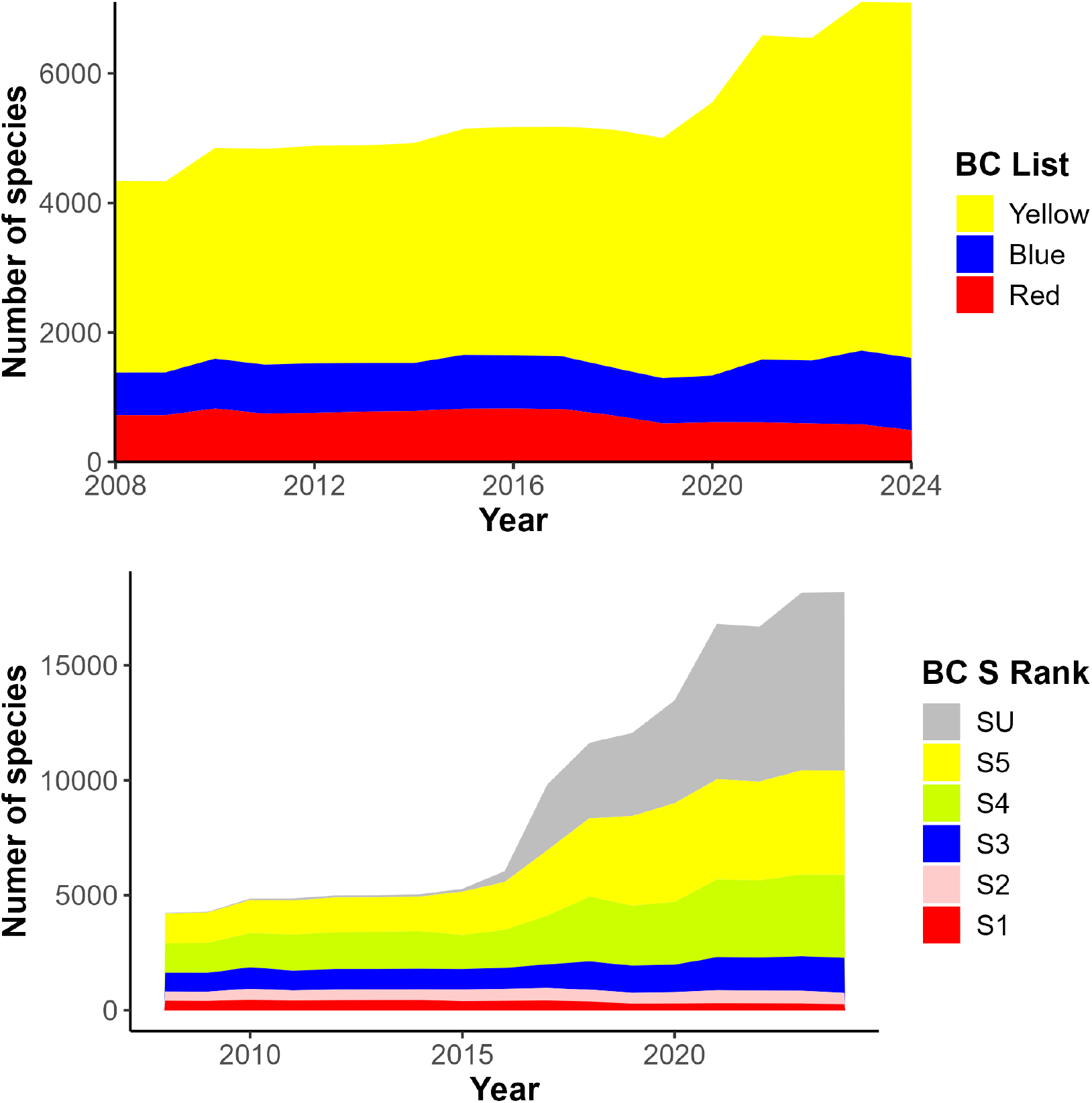
Changes in number of species by B.C. List category (top panel) and NatureServe S Rank (bottom panel). The large jump in size of the Yellow List between 2019-2021 was due to the expansion of the database to include thousands of microlichen and macrofungi (Government of British Columbia 2020), while declines on the Red List around this time reflect non-genuine adjustments in the status of plants (Government of British Columbia 2019).

**Figure 2:**
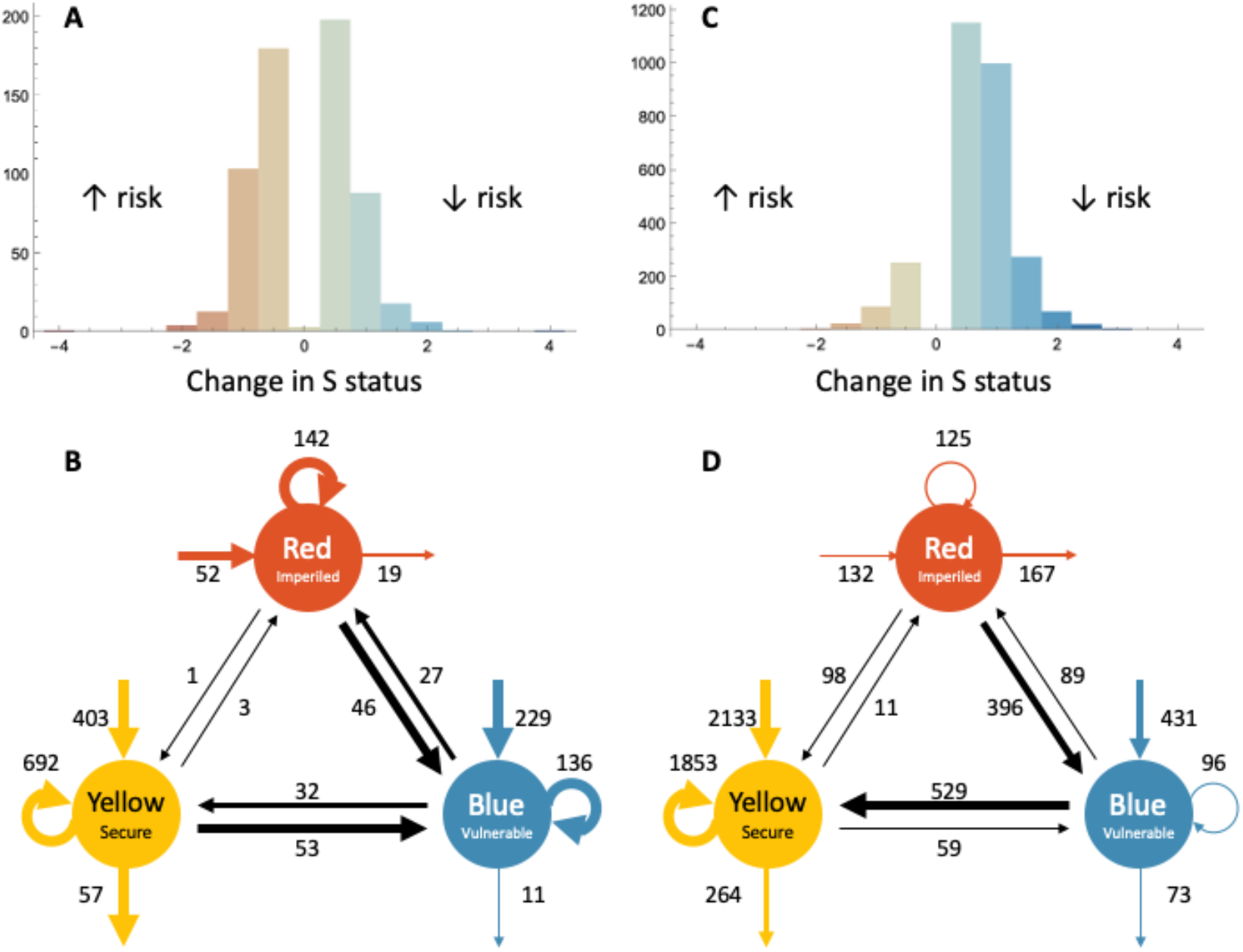
Changes over time in the status of wildlife in British Columbia. Changes in S rank (top) and B.C. List status (bottom) for animals (A,B) and plants (C,D). For the bottom panels, numbers give the total that moved between risk categories based on the annual change reports from 2008-2024, whether the reasons for the change were genuine or not. Line widths are proportional, up to a maximum width of 6pt for (B) 60 animal species or (D) 600 plant species. Species leave the B.C. List for a number of reasons, including that they were never regularly present in B.C., they were combined with another taxonomic unit, or they were split into two units that each have their own status.

The total number of species at risk (Red- or Blue-listed) has increased by 16.3% since 2008, from 1,382 (*n*=658 Blue List, *n*=724 Red List) to 1,607 in 2024. By contrast, the total number of Red-listed species decreased from 2008 to 2024 (from 724 to 491). Many of the species taken off of the Red List were plants and moved for non-genuine reasons, as discussed in the next sub-section (e.g., to align with Canada-wide assessments, see Government of British Columbia, 2019).

### Changes in Status

Between 2008 and 2024, there were 967 changes to the S rank of animal species (Figure 1A). Nearly equal numbers of S rank changes involved improvements to the status of species (“downlisted”, *n*=315) and deteriorating status (“uplisted”, *n*=302). Similarly, roughly equal numbers of changes to the colour-coded B.C. List were downlisted (*n*=79) and uplisted (*n*=83) (Figure 1B). Focusing on changes to the B.C. List that evaluators unanimously interpreted as genuine, animal species became more or less endangered at similar rates (Table 1). Based on the textual comments, slightly more explanations involved genuine changes to the status for uplisted species (27.5%, *n*=19/69) than for downlisted species (18.8%, *n*=12/64; *p*=0.31 Fisher’s exact test). The rank change reasons, which were more reliably interpreted, revealed a similar pattern, with slightly more explanations involving genuine changes for uplisted species (32.4%, *n*=11/34) than for downlisted species (28.1%, *n*=9/32; *p*=0.79 Fisher’s exact test; see list of species in Table 2). Although not statistically significant, these trends suggest that a change in status was more likely to reflect genuine changes when an animal became more endangered than when it became less endangered in B.C. (more often for the comments by 50% and for the reasons by 15%).

**Table 2:**
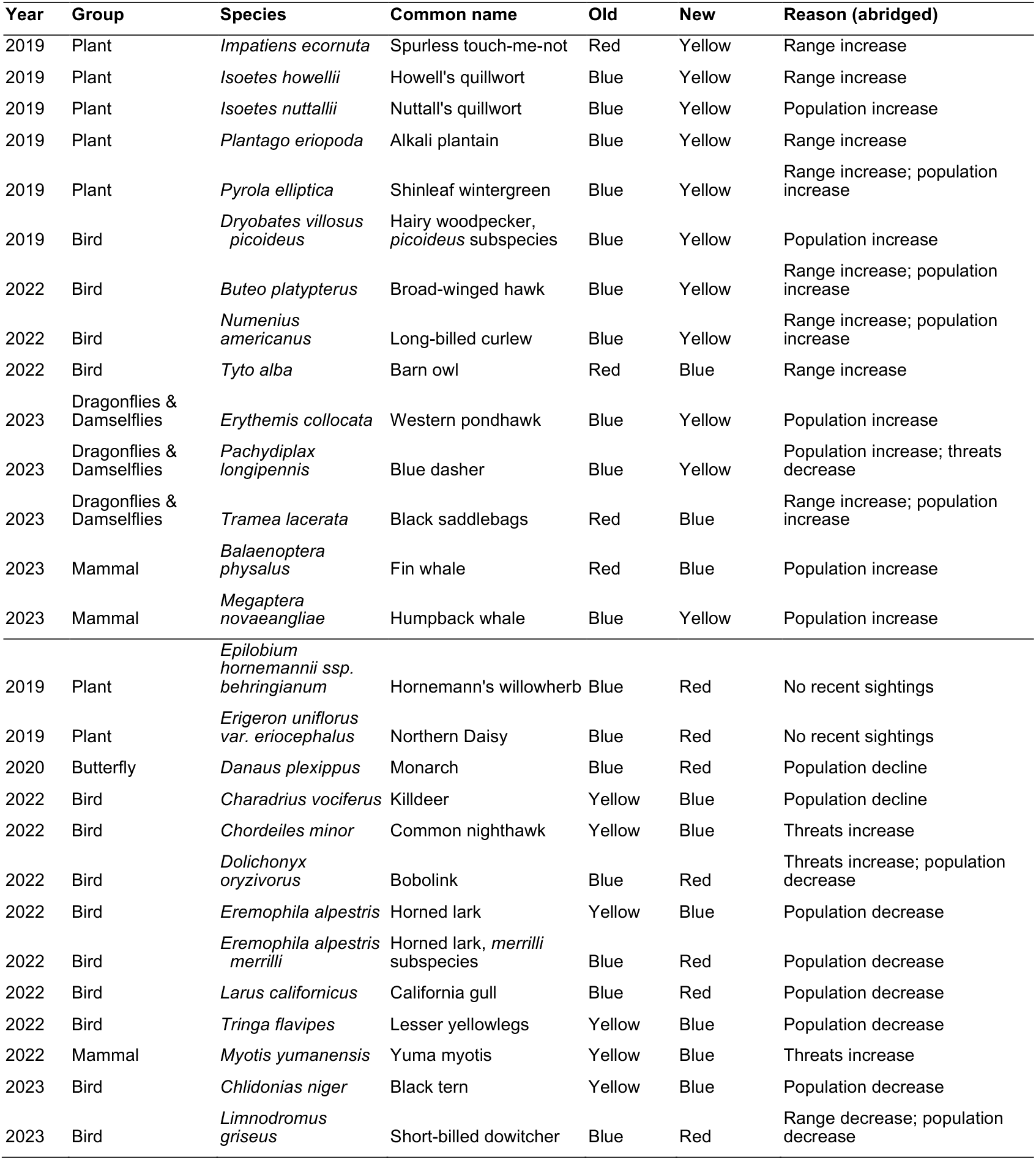
Telling their stories. Species with genuine reasons for a change in B.C. List status. The year of the status change, as well as the old and new colour code, are given for 9 animal and 5 plant downlisted species (top) and 11 animal and 2 plant uplisted species (bottom). Only “reasons” interpreted unanimously by the evaluators are included (2019-2024).

Plants exhibited S rank changes much more frequently than animals (*n*=2,902; Figure 1C). The vast majority of these S rank changes were downlisted (*n*=2,521) rather than uplisted (*n*=381). Similarly, the B.C. List status of plant species improved (*n*=2,023 downlisted) much more often deteriorated (*n*=159 uplisted; Figure 1D). While these changes to plant taxa are superficially encouraging, the vast majority of changes in B.C. List status were non-genuine (Table 1), with only 3% of reasons and 2% of comments evaluated as genuine unanimously among all three authors. Where there was unanimity in the evaluations, uplisting of plants to a more endangered status on the B.C. List was more often deemed genuine (17.0%, *n*=9/53) than downlisting (3.4%, *n*=9/264; *p*=0.0007 Fisher’s exact test). Based on the reasons (excluding 2018), which were more reliably interpreted, uplisting was again more often deemed genuine (8.3%, *n*=2/24) than downlisting (1.4%, *n*=5/366; *p*=0.063 Fisher’s exact test). Overall, the proportion of species that changed in status for genuine reasons was 5 to 6 times higher for plant species that became more endangered than for those that became less endangered.

On average, NatureServe S ranks for species in B.C. became more uncertain over time, with increasing use of ranks such as “?”, “U”, or “Unknown” (animals: 180 more uncertain versus 51 less uncertain; plants: 559 more uncertain versus 160 less uncertain), a point we return to in the Discussion.

### Provincial versus Federal Status

Of the species assessed by the B.C. CDC (February 10, 2025), 370 have also been assessed at the federal level by COSEWIC. The majority of the 221 B.C. species assessed by COSEWIC as either “Endangered” or “Threatened” were also assigned a provincial S Rank of similar intensity (*n*=168 species; 76.0%; equivalent to being Red listed). The same applies for the 84 species assessed as “Special Concern” at the federal level (*n*=52 Blue-listed species; 61.9%) as well as the 69 species assessed as “Not at Risk” by COSEWIC (*n*=46 Yellow-listed species; 66.7%). The majority of species in B.C. with high S Ranks have not, however, been assessed by COSEWIC, reflecting, in part, a backlog in assessments at the federal level (OAG, 2024).

Conversely, some species in B.C. have been recommended for federal listing by COSEWIC (13 Special Concern, 5 Threatened, 2 Endangered) despite being left off of the Blue and Red Lists in B.C. For example, the Alkaline Wing-nerved Moss (*Pterygoneurum kozlovii*) has most recently been assessed by COSEWIC as Threatened, yet its S Rank is S3S4, and it is consequently Yellow-listed at the provincial level. This discrepancy is notable because apart from a small and unconfirmed population in Saskatchewan, all observations of the moss in Canada come from British Columbia (COSEWIC, 2004).

Most imperilled species did not change status at the federal or provincial level during our 15-year study period (Table 3). Specifically, 305 of the 370 species with COSEWIC and provincial assessments in 2024 also had both assessments in 2008, and 230 of those species (75.4%) had the same federal and provincial statuses in both those years. Species that were re-assessed and either uplisted or downlisted by COSEWIC at any time between 2008 and 2024 underwent similar listing changes at the provincial level 35.5% of the time (11 concordant changes, 1 uplisted and 10 downlisted, out of 31 species with COSEWIC listing changes). For nine of these 11 cases, the COSEWIC listing change came before the provincial listing change, with a time lag between the changes under four years for all but one case.

**Table 3:**
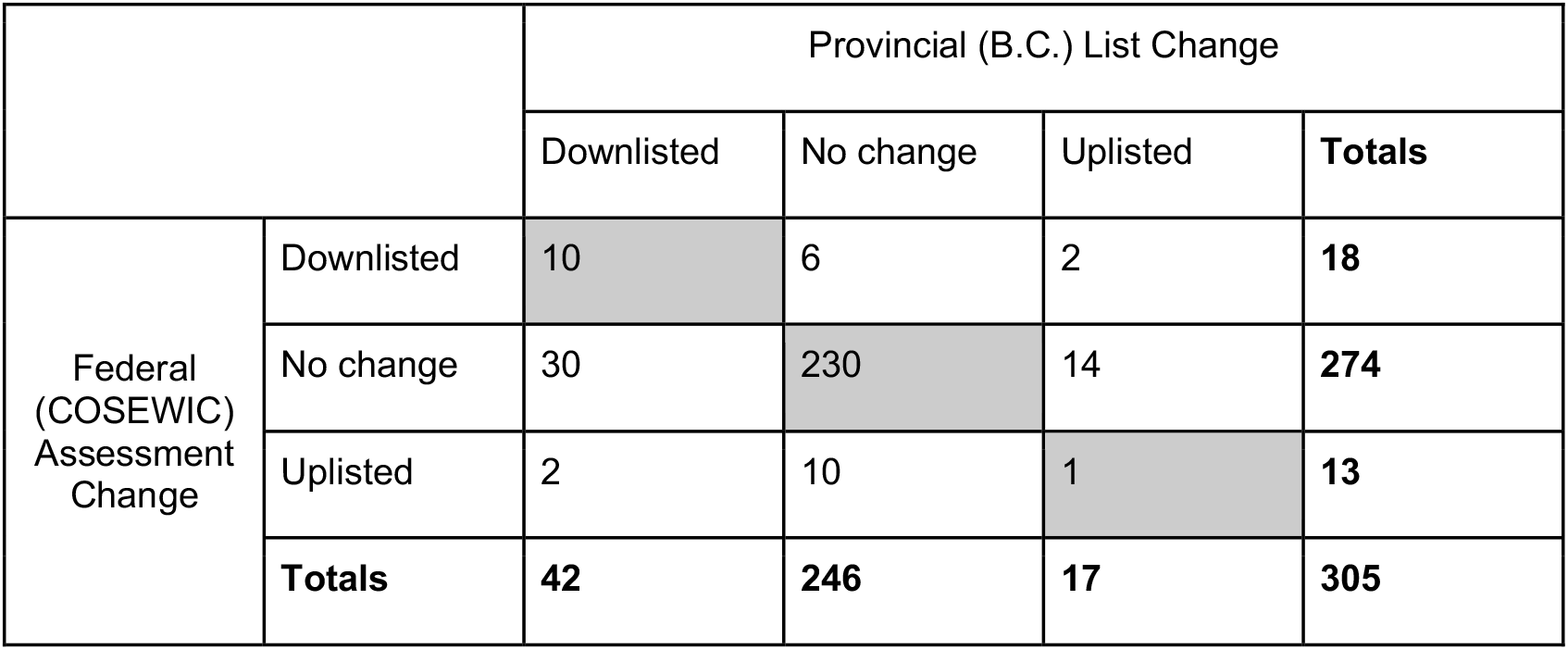
Comparing federal and provincial assessments of species at risk in Canada. Only species that were assessed by COSEWIC and on the B.C. List in both 2008 and 2024 are included. The numbers of uplisted species (increase in endangerment), species with no change in status, and downlisted species (decrease in endangerment) over this time period are shown. The two species that were uplisted federally but downlisted provincially are the Long-billed Curlew (*Numenius americanus*) and Lewis’s Woodpecker (*Melanerpes lewis*), while Bobolink (*Dolichonyx oryzivorus*) and Common Nighthawk (*Chordiles minor*) were uplisted provincially and downlisted federally.

#### Case Studies

To highlight the complex status of B.C.’s endangered species, we present four case studies: the first highlights genuine improvements in provincial status that led to downlisting on the B.C. List; the second is a species that is considered to be more at risk in B.C. according to the federal assessment than the provincial assessment; the third is an example where status changes have gone in opposite directions at the federal and provincial levels; and the final example features a “ghost” species that is ranked as critically imperiled (S1) but is not listed provincially or federally.

Three dragonfly species, the Blue Dasher (*Pachydiplax longipennis*), Western Pondhawk (*Erythemis collocata*), and Black Saddlebags (*Tramea lacerata*), have all undergone range expansions and population increases in B.C. according to a CDC reassessment in 2023. All three species have been observed more frequently in Canada in recent years, independent of observer effort, likely confirming the increase in range, with warming temperatures appearing to be the primary driver of range expansion. For example, the Black Saddlebags was first detected in B.C. in 1996 (Cannings, 1997), and northward range shifts in this species’ distribution across the continent have been attributed to global climate change (Makepeace & Lewis, 2016). The increase in abundance of these dragonfly species in B.C. led to downlisting each species (from Blue to Yellow for *P. longipennis* and *E. collocata*; from Red to Blue for *T. lacerata*), therefore representing genuine improvements in these species’ B.C. status. None of these species have been assessed at the federal level by COSEWIC.

The western rattlesnake (*Crotalus oreganus*) is distributed throughout western North America; however within Canada, it is only found in southern interior B.C. The rattlesnake has a narrow range within B.C. and is located within the Thompson-Okanagan interior east of the Cascade Mountains (Kirk et al., 2021). It is B.C.’s only venomous snake, and consumes a diet of mostly rodents, many of which are considered agricultural pests. The rattlesnake requires two main types of habitat: dens on south facing slopes in fractured rock outcrops that remain above freezing throughout the winter and are thus suitable as winter hibernacula, and summer foraging habitat that consists of areas with cover such as rocks or vegetation (e.g., brush) (COSEWIC, 2015). Threats to the western rattlesnake include habitat loss and fragmentation via urbanization and agriculture (Lomas et al., 2019). The rattlesnake was first designated as Threatened by COSEWIC in 2004 and subsequently reassessed and confirmed in 2015 (COSEWIC, 2015). Although the provincial equivalent to this designation is the Red List, the rattlesnake remains on the Blue List in B.C. (equivalent to “Special Concern” at the federal level). Their official B.C. ranking changed from S3 to S2S3 in 2018 and the associated comment indicated that the change was non-genuine (“This is not a genuine change in status, but a refinement of the previous rank calculator scores based on data from COSEWIC”). The rattlesnake thus serves as an example where the federal process evaluates the extinction risk of the species to be higher than does the provincial process, even based on the same data.

The Bobolink (*Dolichonyx oryzivorus*) is a medium-sized migratory songbird that breeds across southern Canada in various grassland habitats. Breeding habitat for grassland birds is disappearing rapidly across Canada (Daniel and Koper, 2019), including in B.C. (Haddow et al., 2013). In 2022, the Bobolink was moved to a more imperilled status on the B.C. List, changing from Blue (S3) to Red (S2?). The B.C. CDC indicated that this change was genuine based on population declines and increasing threats. By contrast, COSEWIC reassessed the species in the same year (2022) and recommended that it be downlisted from “Threatened” to “Special Concern”, despite a high overall threat and ongoing rates of decline of 25% per decade, because this rate of decline was too slow to merit “Threatened” status. Although currently listed under *SARA* as Threatened, if this change in status is accepted, the Bobolink may soon lose most of its legal protection federally. The B.C. breeding population of Bobolinks is spatially disconnected from the rest of the Canadian population and, with decreasing federal protections on the horizon, stronger provincial protections would be required to reverse declines of Bobolink, and other grassland species, in B.C.

Globally, spruce dwarf tarantulas (*Hexura picea*) are found only along the Pacific coast of North America, including in B.C. and as far south as Northern California (Opler and Lattin, 2001). These tarantulas inhabit cool and moist areas within primarily old growth forests, which have been decreasing throughout its range (Price et al., 2021). Relative to other spider species, they have limited ability to disperse as juveniles (Bond and Godwin, 2013), which could be detrimental in the face of habitat alteration. On iNaturalist, there have been 28 total observations, with only five in Canada (iNaturalist, 2025), and they are generally regarded as rare (Humble et al., 1999). The spruce dwarf tarantula has not been assessed by COSEWIC and is thus not currently listed federally under *SARA*. Provincially, it is a ghost species, with a NatureServe rank of “critically endangered” within B.C. (S1) and Canada (N1), which should place it on the Red List. Given its extremely limited range and threats associated with habitat alteration, including to old growth forests, the spruce dwarf tarantula is an endangered species lacking any legal protection in Canada.

## DISCUSSION

Trends in population size and range are important indicators of a nation’s alignment with global biodiversity targets (e.g., Kunming-Montreal Global Biodiversity Framework Target 4). In B.C., Canada, a three-tiered, colour-listing system categorizes >20,000 species found in the province into categories (Red, Blue, Yellow) corresponding to their population trends, provincial distribution, and risk of extinction. To assess whether B.C. has been effectively managing its biodiversity, we collated and analyzed data from the B.C. Conservation Data Centre (2025) database, dating back to 2008.

At first glance, the trends are promising, with declines in the number of species on the Red List (designated as most endangered) over time (Figure 1). However, through a consensus-based process, we determined that most of these changes in species status were not genuine. In particular, hundreds of plant species were moved off of the Red List due to methodological changes, without a genuine improvement in the population size or range of the species or a reduction in threats faced. Even with the non-genuine downlisting of hundreds of B.C. plant species, we found that the total number of species at risk in B.C. (Red and Blue Lists) has increased by 16.3% since 2008. Overall, very few species have undergone a genuine improvement in status (Table 1). Similar results were reported by Teucher and Ramsay (2013), who back-corrected the data in cases of non-genuine changes and reported that birds, mammals, and reptiles/amphibians became slightly more imperilled in B.C. between 1992-2012, with fish exhibiting little overall change. These results echo Favaro et al. (2014), who noted the lack of progress in protecting Canada’s most at risk species at the federal level, with many species facing the same risks as they did in previous assessments. Indeed, using COSEWIC’s last two assessments, Environment and Climate Change Canada (2025) reports that 80% of species when reassessed retain the same (64%) or worse (16%) status. As argued by Favaro et al. (2014), a lack of change in extinction risk should not be interpreted as a conservation victory. Despite Canada’s commitment to the Kunming-Montreal Global Biodiversity Framework calls for urgent action “for the recovery and conservation of species, in particular threatened species, to significantly reduce extinction risk” (Target 4), the most common outcome for at-risk species in B.C. and in Canada is to remain at a high level of endangerment for decades.

Manually classifying all B.C. listing changes from the past 15 years as “genuine” or “non-genuine” allowed us to uncover important information about the overall directionality of biodiversity trends in B.C. Focusing on the more reliable field (the “reasons” for change), 70% of animal species and 85% of plant species that changed in colour-status on the B.C. List involved non-genuine changes (Table 1). A higher proportion of the reasons were “genuine” for species that became more endangered (11/34 = 32.4% for animals; 2/24 = 8.3% for plants) than for species that became less endangered (9/32 = 28.1% for animals; 5/366 = 1.4% for plants). Because the majority of listing changes were “non-genuine” across all taxonomic groups and directions of change, reports on the status of at-risk wildlife populations over time will artificially paint a rosier picture of biodiversity trends in B.C. This echoes concerns raised at the federal level that improvements in status of species in Canada often involve non-genuine changes to our knowledge, while declines in species’ status were typically genuine (Moore et al., 2016). It is important that the status of species is updated in light of emerging information, and these non-genuine changes to the record are important to avoid misplaced conservation efforts. Nevertheless, it is the genuine changes that inform us about the efficacy of efforts to protect species at risk.

We found that genuine changes in extinction risk of B.C. species nearly equally often involved increases in risk as decreases in risk. Considering comments and reasons from CDC annual change reports that were unanimously deemed genuine or non-genuine, we found that genuine uplistings outnumbered genuine downlistings (28 versus 21) according to the comments and were slightly exceeded by genuine downlistings (13 versus 14) according to the reasons. These numbers reinforce the message that genuine improvements in the status of species at risk have been exceedingly rare in B.C. These findings highlight the limited scope of the Wildlife Act, which only provides legal protection to four species, leaving many others in jeopardy. More robust protections and associated conservation efforts could have led to more genuine uplistings; however, this was not the case, highlighting the deficiency of biodiversity protections in B.C. (Westwood et al., 2019).

Despite the expectation that biodiversity knowledge would increase over time, we found that the status of species became 3 to 4 times more uncertain following a change in “S Rank” in British Columbia, with newer ranks being more likely to involve “?” or “U” (unknown). This increase in uncertainty over time may reflect shifts in how the existing uncertainty is recorded, although it is possible that threats are becoming increasingly challenging to quantify. Additional uncertainty could be facing species subjected to new threats (e.g., newly introduced species). Existing information on species status also becomes more outdated over time, increasing uncertainty in the absence of new data (Kindsvater et al., 2018). Alternatively, the scientific community may be more aware of data limitations, with changes in status better accounting now for uncertainty that has always existed. The addition of over 10,000 species to the database over our 15-year study period may also skew the species pool to understudied or poorly understood species for which status estimates would likely be uncertain. Regardless, our analysis emphasizes the importance of continuing to collect, analyze, and review data on biodiversity in B.C., as reducing uncertainty in S Ranks will help bring conservation attention to all imperiled species and help allocate conservation resources appropriately.

The data managed by the B.C. CDC provide an invaluable snapshot of the status of monitored species in the province. Recent changes to the database have improved its utility, including the addition of a “rank change reason” with clearer wording about which changes are genuine. As of this year, previous snapshots of the database are now also available back to 2011 (Government of British Columbia 2025). Nevertheless, there is a great deal of uncertainty in the status of species in B.C. (Figure 1B). Over 1000 “ghost” species have received NatureServe rankings indicating that they are at risk in B.C., but these remain to be officially reviewed and placed on the B.C. List of Red and Blue species. As a consequence, there is considerable doubt as to whether wildlife populations are improving or declining in B.C. and how many species are actually at risk without more investment in monitoring biodiversity within the province.

This study underscores the unfortunate truth that even the Red- and Blue-listed species in British Columbia are not afforded effective legal protections by the province despite significant efforts to assess thousands of species and populations. The “S Rank” system designed by NatureServe conveys both the intensity and uncertainty of a species’ extinction risk and translates directly to the colour-coded listing scheme adopted by the province. However, without any formal legislation to protect these species and their habitats, these listings are no more than an admission that highly endangered species exist in B.C. and that the vast majority of species that were at risk 15 years ago remain at risk. Canada’s federal *SARA* protects species and their habitats if these species are deemed to be sufficiently at risk of extinction at the federal level, but this act comes with caveats: 1) most protective powers only cover federal lands, which represent only 4% of Canada’s ten provinces (Gordon et al., 2024) and just 1% of B.C.; 2) *SARA* only protects 656 species across Canada and 256 in B.C., a small fraction of the 2,614 species at risk in the province (1,116 Blue-Listed, 491 Red-Listed, and 1,012 “ghost” species with a NatureServe status meriting Red- or Blue-listing), with a significant bias toward larger animal species; and 3) designatable units may be at risk in BC but stable across the rest of their range so are not eligible for *SARA* listing, which could have consequences for regional biodiversity. Even with federal protections in place for Canada’s wildlife, provincial protections are needed to fill important gaps and conserve species where they actually live. To support biodiversity conservation and address the gap in protections for listed species at risk in B.C., a dedicated and effective provincial species at risk law is needed. Our results show minimal genuine changes to status ranks over the last 15 years in B.C., indicating that the state of biodiversity within the province is not improving and that the existing legal and regulatory mechanisms are not sufficient to protect or recover at-risk species in B.C.

## ACKNOWLEDGEMENTS

We thank the B.C. Conservation Data Centre for their work to assess species in the province and for providing the data used in this study. We would like to thank Dr. Arne Mooers for reviewing an early draft of this article and the Canadian Parks and Wilderness Society, B.C. Chapter (CPAWS-B.C.) Land and Freshwater Team for their support of this project, including salary for MB during the early stages.

## Data sources

Historical records for species at risk from the B.C. CDC are available for download for the years 2011-2024 from: https://catalogue.data.gov.bc.ca/dataset/bc-species-and-ecosystems-historical-conservation-status-information/resource/c7247ad5-3625-433c-8355-ac7264656731

Data from 2008-2010 were kindly provided by the B.C. Conservation Data Centre.

Information about changes to the status and name of species in B.C. are posted by year and were downloaded from the CDC (https://www2.gov.bc.ca/gov/content/environment/plants-animals-ecosystems/conservation-data-centre/explore-cdc-data/conservation-data-centre-updates; accessed 6 March 2025).

Data and code necessary to reproduce our analyses are available at the following GitHub repository: https://github.com/pthompson234/bc-species-at-risk

## Notes

https://github.com/pthompson234/bc-species-at-risk

### Competing Interest Statement

Competing interests: Meg Bjordal was an employee of the Canadian Parks and Wilderness Society, BC Chapter during the early stages of this project. The other authors declare no competing interests.

### Funding

- Natural Sciences and Engineering Research Council: RGPIN-2022-03726
- L'Oreal (United States): Award For Women in Science
- Liber Ero Fellowship: Post-doctoral Fellowship
- Canadian Parks and Wilderness Society, BC Chapter: Salary support (partial for MB)

## REFERENCES

B.C. Conservation Data Centre. 2025. BC Species and Ecosystems Explorer. B.C. Government, Victoria B.C. Available: http://a100.gov.bc.ca/pub/eswp/

B.C. Conservation Data Centre. 2023. Red, Blue & Yellow Lists. Available from: https://www2.gov.bc.ca/gov/content/environment/plants-animals-ecosystems/conservation-data-centre/explore-cdc-data/red-blue-yellow-lists. Accessed June 3, 2025.

Bolliger CS, Raymond CV, Schuster R, and Bennett JR. 2020. Spatial coverage of protection for terrestrial species under the Canadian Species at Risk Act. Écoscience, 27(2): 141–147. 10.1080/11956860.2020.1741497

Bond JE and Godwin RL. 2013. Taxonomic revision of the trapdoor spider genus Eucteniza Ausserer (Araneae, Mygalomorphae, Euctenizidae). ZooKeys, 356: 31. 10.3897/zookeys.356.6227

Cannings RA. 1997. Tramea lacerata (Hag.) new to British Columbia, Canada, with notes on its status in the northwestern United States (Anisoptera: Libellulidae). Notulae Odonatologicae, 4(9): 148–149. https://natuurtijdschriften.nl/pub/593594

Catling PM. 2016. Climate warming as an explanation for the recent northward range extension of two dragonflies, Pachydiplax longipennis and Perithemis tenera, into the Ottawa Valley, eastern Ontario. The Canadian Field-Naturalist, 130(2): 122–132. 10.22621/cfn.v130i2.1846

COSEWIC. 2004. COSEWIC assessment and status report on the alkaline wing-nerved moss Pterygoneurum kozlovii in Canada. Committee on the Status of Endangered Wildlife in Canada Ottawa. vi + 20 pp. (www.sararegistry.gc.ca/status/status_e.cfm).

COSEWIC. 2015. COSEWIC assessment and status report on the Western Rattlesnake Crotalus oreganus in Canada. Committee on the Status of Endangered Wildlife in Canada. Ottawa. xi + 44 pp. (Species at Risk Public Registry).

Daniel J and Koper N. 2019. Cumulative impacts of roads and energy infrastructure on grassland songbirds. The Condor: Ornithogical Applications, 121(2): duz011. 10.1093/condor/duz011

Environment and Climate Change Canada. (2025). Canadian Environmental Sustainability Indicators: Changes in the status of wildlife species at risk. https://www.canada.ca/en/environment-climate-change/services/environmental-indicators/changes-status-wildlife-species-risk.html.

Faber-Langendoen D, Nichols J, Master L, Snow K, Tomaino A, Bittman R, Hammerson G, Heidel B, Ramsay L, Teucher A, and Young B. 2012. NatureServe Conservation Status Assessments: Methodology for Assigning Ranks. NatureServe, Arlington, VA.

Favaro B, Claar DC, Fox CH, Freshwater C, Holden JJ, Roberts A, and Derby UVR. 2014. Trends in extinction risk for imperiled species in Canada. PLOS ONE, 9(11): e113118. 10.1371/journal.pone.0113118

Forest Practices Board (FPB). 2023. Management of habitat for species at risk under the Forest and Range Practices Act. Forest Practices Board, Victoria, B.C. Special Investigation No. 55. 24 p.

Gordon SCC, Duchesne AG, Dusevic MR, Galán-Acedo C, Haddaway L, Meister S, Olive A, Warren M, Vincent JG, Cooke SJ, and Bennett JR. 2024. Assessing species at risk legislation across Canadian provinces and territories. FACETS, 9: 1–18. 10.1139/facets-2023-0229

Government of British Columbia. 2019. BC Conservation Status Rank Review and Changes (Vascular plants and macrolichens). Accessed July 21, 2025, from https://www2.gov.bc.ca/assets/gov/environment/plants-animals-and-ecosystems/conservation-data-centre/data-changes/2019_plant_lichen_rank_review_changes.pdf

Government of British Columbia. 2020. BC Conservation Status Rank Review and Changes (Vascular and nonvascular plants, lichens and macrofungi). Accessed July 21, 2025, from https://www2.gov.bc.ca/assets/gov/environment/plants-animals-and-ecosystems/conservation-data-centre/data-changes/2020_plant_fungi_changes_summary.pdf

Government of British Columbia. 2025. BC Species and Ecosystems Previous Years Conservation Status - Compilation https://catalogue.data.gov.bc.ca/dataset/bc-species-and-ecosystems-previous-years-conservation-status-compilation

Government of Canada. 2025. Species at risk public registry [online]: Available from canada.ca/en/environment-climate-change/services/species-risk-public-registry.html.

Greenwald N, Suckling KF, Hartl B, and Mehrhoff LA. 2019. Extinction and the US Endangered Species Act. PeerJ, 7: e6803. 10.7717/peerj.6803

Haddow C, Bings B, and Wallich E. 2013. Cover Requirements and Habitat Needs of Grassland-nesting Birds in the Cariboo-Chilcotin. Ministry of Forests, Lands and Natural Resource Operations, Resource Practices Branch, Victoria, B.C. FREP Report. 36 p.

Humble LM, Winchester NN, and Ring RA. 1999. The potentially rare and endangered terrestrial arthropods in British Columbia: revisiting British Columbia’s biodiversity. In Proceedings of a Conference on the Biology and Management of Species and Habitats at Risk, Kamloops, B.C., 15–19 February 1999. pp. 15–19.

iNaturalist. 2025. Available from: https://www.inaturalist.org/taxa/172579-Hexura-picea#cite_note-:1-6. Accessed April 24, 2025.

International Union for Conservation of Nature and Natural Resources. 2025. Reasons for changing category. The IUCN Red List of Threatened Species. Available from: https://www.iucnredlist.org/assessment/reasons-changing-category. Accessed July 21, 2025.

Kindsvater HK, Dulvy NK, Horswill C, Juan-Jordá MJ, Mangel M, Matthiopoulos J. 2018. Overcoming the data crisis in biodiversity conservation. Trends in Ecology & Evolution. 33(9):676–88.

Kirk DA, Karimi S, Maida JR, Harvey JA, Larsen KW, and Bishop CA. 2021. Using ecological niche models for population and range estimates of a threatened snake species (Crotalus oreganus) in Canada. Diversity, 13(10): 467. 10.3390/d13100467

Lomas E, Maida JR, Bishop CA, and Larsen KW. 2019. Movement ecology of northern Pacific rattlesnakes (Crotalus o. oreganus) in response to disturbance. Herpetologica, 75(2): 153–161. 10.1655/D-17-00060

Mace GM, Reyers B, Alkemade R, Biggs R, Chapin FS, Cornell SE, Díaz S, Jennings S, Leadley P, Mumby PJ, Purvis A, Scholes RJ, Seddon AWR, Solan M, Steffen W, and Woodward G. 2014. Approaches to defining a planetary boundary for biodiversity. Global Environmental Change, 28: 289–297. 10.1016/j.gloenvcha.2014.07.009

Makepeace HS and Lewis JH. 2020. New and notable records of Odonata from New Brunswick, Canada, with a significant eastern range extension of Enallagma anna. Northeastern Naturalist, 27(4): N58. 10.1656/045.027.0406

Moore A, Cyr A, and Findlay S. 2016. Do changes in COSEWIC status reflect changes in species’ biological status. Report to the Committee on the Status of Endangered Wildlife in Canada (COSEWIC), 10 pp.

NatureServe (2025) Rank change reason guidance. Available from: https://help.natureserve.org/biotics/content/record_management/Element_Files/Element_Tracking/Rank_Change_Reason_Guidance.htm. Accessed September 27, 2025.

Office of the Auditor General of British Columbia (OAG B.C.). 2021. Management of the conservation lands program. Office of the Auditor General of British Columbia. https://www.oag.bc.ca/app/uploads/sites/963/2024/08/OAGB.C.-20210511-Conservation-Lands-Program_RPT.pdf

Office of the Auditor General of Canada (OAG). 2024. Supporting Species at Risk Assessment and Reassessment—Environment and Climate Change Canada (Independent Auditor’s Report No. 9; Reports of the Commissioner of the Environment and Sustainable Development to the Parliament of Canada). Office of the Auditor General of Canada. https://www.oag-bvg.gc.ca/internet/docs/parl_cesd_202411_09_e.pdf

Opler PA and Lattin JD. 2001. Systematic Compendium. In: Narrative on arthropods and annelid worms of old-growth and late successional conifer forests, mature riparian woods, and of coarse woody debris associated arthropods within the range of the northern spotted owl (Strix occidentalis caurina). Available from: http://www.mesc.usgs.gov/resources/education/arthropods/systematic_compendium.html

Price K, Holt RF, and Daust D. 2021. Conflicting portrayals of remaining old growth: the British Columbia case. Canadian Journal of Forest Research, 51(5): 742–752. 10.1139/cjfr-2020-0453

Province of British Columbia. 2023. Together for Wildlife - Action 10: Spatial Analysis of Disturbance within Habitat Designations in British Columbia. Ministry of Water, Lands, and Resource Stewardship and Ministry of Forests, Together for Wildlife Action 10 Team, Victoria, B.C. FREP Report No. 45. 52 p.

Teucher A and Ramsay L (2013) Trends in the status of native vertebrate species in B.C. Accessible at: https://www2.gov.bc.ca/assets/gov/environment/research-monitoring-and-reporting/reporting/envreportbc/archived-indicators/plants-and-animals/2013_trends_native_vertebrate_species_1992-2012.pdf, with methods at: https://www.env.gov.bc.ca/soe/indicators/plants-and-animals/print_ver/2013_Trends_Native_Vertebrates_Methods.pdf.

Toochin R. 2024. B.C. rare bird record (January 31 2024 entry). https://bcrarebirdrecords.ca/species/pinyon-jay

Westwood AR, Otto SP, Mooers A, Darimont C, Hodges KE, Johnson C, Starzomski BM, Burton C, Chan KMA, Festa-Bianchet M, Fluker S, Gulati S, Jacob AL, Kraus D, Martin TG, Palen WJ, Reynolds JD, and Whitton J. 2019. Protecting biodiversity in British Columbia: recommendations for developing species at risk legislation. FACETS, 4(1): 136–160. 10.1139/facets-2018-0042

